# Visual imagery vividness correlates with afterimage conscious perception

**DOI:** 10.1101/2023.12.07.570716

**Authors:** Sharif I. Kronemer, Micah Holness, A. Tyler Morgan, Joshua B. Teves, Javier Gonzalez-Castillo, Daniel A. Handwerker, Peter A. Bandettini

## Abstract

Afterimages are illusory, visual conscious perceptions. A widely accepted theory is that afterimages are caused by retinal signaling that continues after the physical disappearance of a light stimulus. However, afterimages have been reported without preceding visual, sensory stimulation (e.g., conditioned afterimages and afterimages induced by illusory vision). These observations suggest the role of top-down, brain mechanisms in afterimage conscious perception. Therefore, some afterimages may share perceptual features with sensory-independent conscious perceptions (e.g., imagery, hallucinations, and dreams) that occur without bottom-up, sensory input. In the current investigation, we tested for a link between the vividness of visual imagery and afterimage conscious perception. Participants reported their vividness of visual imagery and perceived sharpness, contrast, and duration of negative afterimages. The afterimage perceptual features were acquired using perception matching paradigms that were validated on image stimuli. Relating these perceptual reports revealed that the vividness of visual imagery positively correlated with afterimage contrast and sharpness. These behavioral results support shared neural mechanisms between visual imagery and afterimages. This study encourages future research combining neurophysiology recording methods and afterimage paradigms to directly examine the neural mechanisms of afterimage conscious perception.

## Introduction

Afterimages are illusory visual perseverations – lasting seconds to minutes – that often follow light stimulation but absent the original inducing light source. Analogous perceptual phenomena are reported in other human senses, including auditory afterimages or *aftersounds*^1–3^. Afterimages have been a source of intrigue for centuries because of their apparent ubiquity, including among non-human animals (e.g., macaques, cats, and pigeons) and its unique insight on the physiological mechanisms of vision^4–7^. In fact, afterimages helped debunk emission theories of vision that explained conscious sight by the projection of light or aether rays from the eyes. Afterimages share perceptual characteristics with aftereffects (e.g., the McCollough effect^8,9^) and filling-in illusions (e.g., Kanizsa or occluded stimuli illusions^10^). However, afterimages are distinct because they do not require concurrent visual input (e.g., they can appear in total darkness).

A widely accepted theory is that afterimages are sourced by retinal signaling following the physical disappearance of a light stimulus. Likewise, afterimages have been described as painted or photographed on the retina^11,12^. Support of afterimage retinal mechanisms includes evidence of fatigue or bleaching of retinal photoreceptors that persistently signal in the absence of physical light stimulation, forming the appearance of an afterimage^7,11,13–18^. Likewise, direct recordings from retinal ganglion cells find a post-receptor rebound response following inducer stimulation that may originate from photoreceptor signaling^19^. A similar retinal-based process is suggested by the opponent-process theory for chromatic afterimages, whereby complementary color afterimages are perceived according to the inducer color (e.g., a yellow inducer forms a blue afterimage), predicted by opponent visual pairs – black-white, red-green, and blue-yellow – so that adaptation to one half of an opponent pair will drive its opposite hue in the subsequent afterimage^11,20^.

However, research also finds that afterimages are not fully explained by retinal mechanisms. This evidence includes that the bleaching of photoreceptors is *not* a necessary condition for the formation of afterimages^6,21,22^. In fact, afterimages can emerge without previous photoreceptor stimulation, as in afterimages by illusory vision (e.g., a perceptually filled-in image)^23^. Likewise, studies find that color spreading in afterimages can extend beyond the boundary of the preceding inducer stimulus^24,25^. There are also reports of afterimages evoked by dreams, imagery, and hallucinations^26–29^. Similarly, conditioned afterimages are reported by pairing tones and inducer stimuli and then withholding the anticipated inducer, yet participants still report seeing afterimages without preceding visual stimulation^30^. These results support that some afterimages are comparable to sensory-independent conscious perceptions (e.g., imagery, hallucinations, and dreams) that are sourced by top-down, brain mechanisms without bottom-up, sensory input.

If some afterimages emerge without retinal signaling, a possible implication is that afterimages are perceptually linked to sensory-independent conscious perceptions that form by top-down, brain mechanisms. Following a similar logic, previous research studied a relationship between the vividness of imagery and the occurrence of hallucinations and dreams^31,32^. While consideration of afterimages as a kind of sensory-independent conscious perception has been discussed, there is limited research on this topic^33^. In fact, there is no previous study comparing the perception of afterimages and imagery. The nearest instances include rare reports of imagery inducing afterimages, afterimages induced by stimuli that were reported as challenging to imagine, and a note by William James (1842-1910) that his visual imagery could be subliminally driven by afterimages^26,34,35^.

To address this gap in the literature, we investigated a possible perceptual link between visual imagery and afterimages. Specifically, we tested if the vividness of visual imagery (i.e., the ability to evoke lifelike visual perception by imagination) correlates with the sharpness, contrast, and duration of afterimages. We hypothesized that if the brain mechanisms involved in visual imagery are shared with those of afterimage conscious perception, then the vividness of visual imagery and afterimages may be linked (e.g., people with more vivid visual imagery also experience more vivid afterimages). Interrogating the perceptual relationship between afterimages and imagery is significant towards resolving the long-standing query for the role of bottom-up, retinal versus top-down, brain mechanisms in afterimage conscious perception^28^.

## Methods

### Participants

Healthy, adult participants (N = 62; males = 22; mean age: 28.90 years; age SD: 10.31 years; mean education = 16.34 years; education SD: 1.91 years) were recruited from the local Bethesda, Maryland community. Two additional participants who completed the study were excluded from analyses because of poor behavioral performance or a corrupted behavioral file. All participants were recruited and consented following protocols approved by the Institutional Review Board of the National Institute of Mental Health. Inclusion criteria included: (1) being between the ages of 18 and 65 years old at the time of experimentation, (2) a healthy physical examine completed by a nurse practitioner within a year of the study session, and (3) ability to give informed consent. Exclusion criteria included: (1) previous or current histories of neurologic or psychiatric disorder, (2) low vision (corrected normal vision was acceptable), and (3) head injuries (e.g., loss of consciousness for >30 minutes and three or more concussive injuries). Prior to each testing session, a nurse practitioner completed a health exam for each participant, including recording temperature, vitals, and assessment for Covid-19 symptoms.

### Afterimage Induction

Afterimages were elicited using an inducer stimulus: a black silhouette image of a human face in frontal view (presentation duration = 4 seconds; visual angle: 4.60 x 8.47 degrees; https://creazilla.com/nodes/2524-face-silhouette; Figure 1C *Inducer Stimulus*; Supplementary Movie 1). The inducer stimulus resulted in negative afterimages that appeared as white or light gray versions of the inducer. In pilot testing, it was observed that some participants perceived an instantaneous, illusory, crisp white version of the inducer stimulus *at the moment* of its disappearance. This experience was sometimes confused with the subsequent negative afterimage that was typically delayed from the offset of the inducer, less sharp than the inducer, and lasted for several seconds. To limit the occurrence of this flashbulb-like perception at the sudden offset of the full contrast inducer, in the first and last second of the inducer presentation, the inducer contrast was gradually ramped up and down to full contrast and no contrast, respectively. Thus, the inducer appeared at full contrast for a total of 2 seconds. In pilot testing (data not shown), the inducer contrast ramping suppressed the perceived offset flash without reducing the occurrence of afterimages.

**Figure 1.**
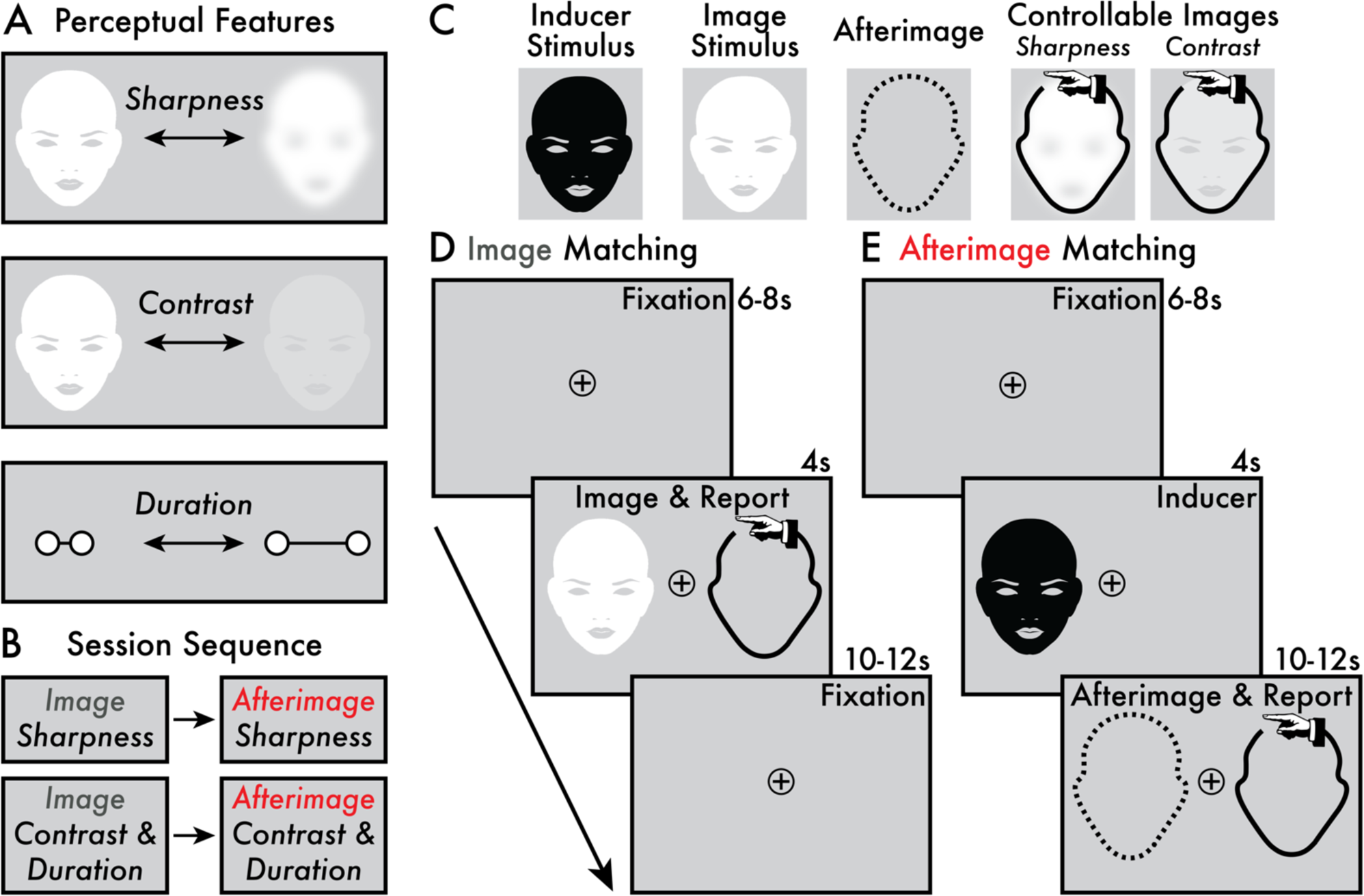
Target perceptual features, session sequence, and perception matching paradigms. **(A)** The target image and afterimage perceptual features were: (1) sharpness, (2) contrast, and (3) duration. **(B)** Participants completed four task phases in the following order: (1) image and (2) afterimage sharpness perception matching and (3) image and (4) afterimage contrast and duration perception matching. **(C)** The stimuli and controllable images presented in the perception matching tasks. The afterimage perception is depicted as a dashed outline because no image was physically presented – the afterimage is an illusory conscious perception. Depending on the task phase, the controllable image allowed participants to manually adjust its sharpness or contrast. The controllable image is depicted with a hand icon (not present during the task) to indicate that participants manually adjusted these images with key presses. **(D)** The main trial phases of the image perception matching task (Supplementary Movies 2 and 4). Each trial began with a fixation interval (6-8 seconds [s]). When the image stimulus appeared (4 s) on either the left or right side of the central fixation, participants were instructed to immediately adjust the controllable image using key presses to match with the image stimuli according to the target perceptual feature (i.e., sharpness and contrast/duration; see *Image Sharpness* and *Contrast and Duration Perception Matching Paradigm* Methods sections). A subsequent fixation interval (10-12 s) followed the *Image & Report* stage prior to initiating the next trial. **(E)** The main trial events of the afterimage perception matching task (Supplementary Movies 3 and 5). Each trial began with a jittered fixation interval (6-8 s). Next, the inducer stimulus was shown (4 s) on either the left or right side of the central fixation and, subsequently, an afterimage might appear. If an afterimage was perceived, participants were instructed to immediately adjust the controllable image to match with the target perceptual feature of their afterimage (i.e., sharpness and contrast/duration; see *Afterimage Sharpness* and *Contrast and Duration Perception Matching Paradigm* Methods sections). The *Afterimage & Report* stage completed when the participant no longer perceived their afterimage, and the remaining duration of time (10-12 s) was a fixation interval prior to initiating the next trial.

During initial task instructions, participants were repeatedly shown the inducer to determine their susceptibility for perceiving afterimages. If there was confusion regarding what parts of their visual experience constituted the afterimage, clarifying instructions were provided by the experimenter to guide when and what parts of their visual perception following the inducer constituted the afterimage conscious perception.

### Image and Afterimage Perceptual Vividness

Participants were asked to report on three target perceptual features that contribute to the overall perceived strength or vividness of conscious vision: (1) *sharpness* (i.e., crisp versus blurry), (2) *contrast* (i.e., bright versus dim), and (3) *duration* (Figure 1A). Sharpness, contrast, and duration are previously interrogated as markers of the vividness of afterimages (e.g., ^17,36^). Here, participants made judgements on these perceptual features for both image and afterimage conscious perception (Figure 1D, E). These perceptual reports were acquired using paradigms where participants adjusted the appearance of an on-screen image – a *controllable image* (Figure 1C *Controllable Images*) – to match in real time with the perceived sharpness, contrast, and duration of images and afterimages. Note that the contrast and duration reports were acquired simultaneously (see *Contrast and Duration Perception Matching* section). The current perception matching approach builds on previous methods for reporting on the perceptual features of afterimages (e.g., ^13,37–39^). Before completing the perception matching tasks, participants were administered instructions and a practice session (see *Sharpness* and *Contrast and Duration Perception Matching* sections). Only when participants felt comfortable with the reporting procedure did they continue to the image and afterimage perception matching tasks (Figure 1B, D, E).

### Sharpness Perception Matching Paradigm

Participants were asked to notice and report on the *maximum* perceived sharpness of images and afterimages. Sharpness was reported using a controllable, on-screen image that participants volitionally manipulated using key presses. See full details below.

#### Image Sharpness Perception Matching

Participants completed an *image* sharpness perception matching task (Figure 1D; Supplementary Movie 2). The image stimulus was a white version of the afterimage inducer stimulus (Figure 1C *Image Stimulus*). The image stimulus appeared for 4 seconds. In the first and last second of presentation, the image stimulus gradually increased and decreased its sharpness, respectively. When the image reached its maximum sharpness, it maintained this value for 2 seconds. This dynamic of increasing and decreasing sharpness was programmed according to pilot testing (data not shown) that suggested this trend matched the perception of afterimage sharpness by the current inducer stimulus.

The sharpness values applied to the image stimulus ranged from 0 (no blurring) to 25 (maximum blurring) in increments of 1, each value representing the number of pixels in the radius of a gaussian kernel used to blur the image stimulus (blurred image size = 600 x 800 pixels; gaussian blur; Illustrator, Adobe, Inc.). Three maximum sharpness values were tested: 10, 15, and 20 pixels. In the analyses and figures (Figure 2C; Figure 3D, E; Figure 4A), the sharpness values were inverted so that 0 pixels indicated the *blurriest* perception and 25 pixels the *sharpest*. Inverting the sharpness pixel scale was implemented because it corresponded with the contrast and duration scales, where larger numbers indicate more vivid images and afterimages. Thus, all sharpness values and accompanying figures are reported along the inverted pixel scale.

**Figure 2.**
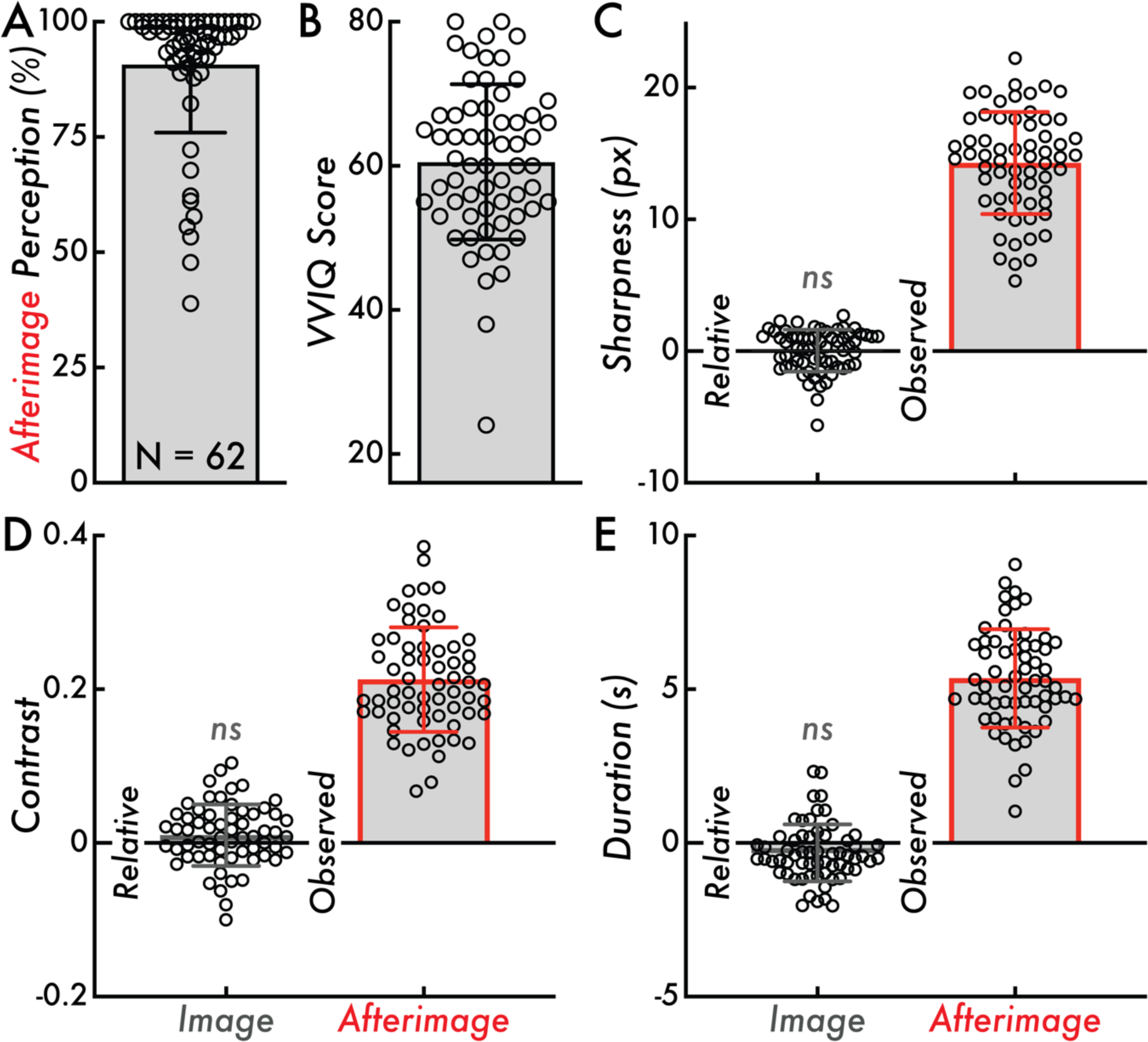
Afterimage perception rate, VVIQ score, and relative image and observed afterimage sharpness, contrast, and duration. **(A)** Afterimage perception rate calculated as the percentage of inducers where a subsequent afterimage was reported across all trials of the afterimage perception matching tasks (90 trials total). The bar graph indicates the mean afterimage perception percentage across participants (90.79%) and the error bar displays standard deviation (SD; 14.84%). **(B)** The Vividness of Visual Imagery Questionnaire (VVIQ) score calculated as the sum of scores across all questionnaire items within participant (score range: 16-80; larger values indicating more vivid visual imagery). The bar graph indicates the mean VVIQ score (60.55) and the error bars display the SD (10.78). **(C)** Relative image and observed afterimage reported maximum sharpness in pixels (px). The relative image sharpness is compared on a trial level against the true image sharpness (true values: 10, 15, or 20 px). The bar height indicates the group mean (*Relative* = 0.033 px; *Observed* = 14.27 px) and the error bars display SD (*Relative* = 1.60 px; *Observed* = 3.88 px). **(D)** Relative image and observed afterimage reported maximum contrast. The relative image contrast is compared against the true image maximum contrast (0.25). The bar height indicates the group mean (*Relative* = 0.01; *Observed* = 0.21) and the error bars display SD (*Relative* = 0.04; *Observed* = 0.068). **(E)** Relative image and observed afterimage reported duration in seconds (s). The relative image contrast is compared against the true image duration (4 s). The bar height indicates the group mean (*Relative* = −0.33 s; *Observed* = 5.35 s) and the error bars display SD (*Relative* = 0.93 s; *Observed* = 1.60 s). Comparing the relative image contrast, sharpness, and duration values from zero was not statistically significant (*ns*; Wilcoxon Rank Sum tests, *p* > 0.05). In all subplots, the open circles represent individual participants (N = 62).

**Figure 3.**
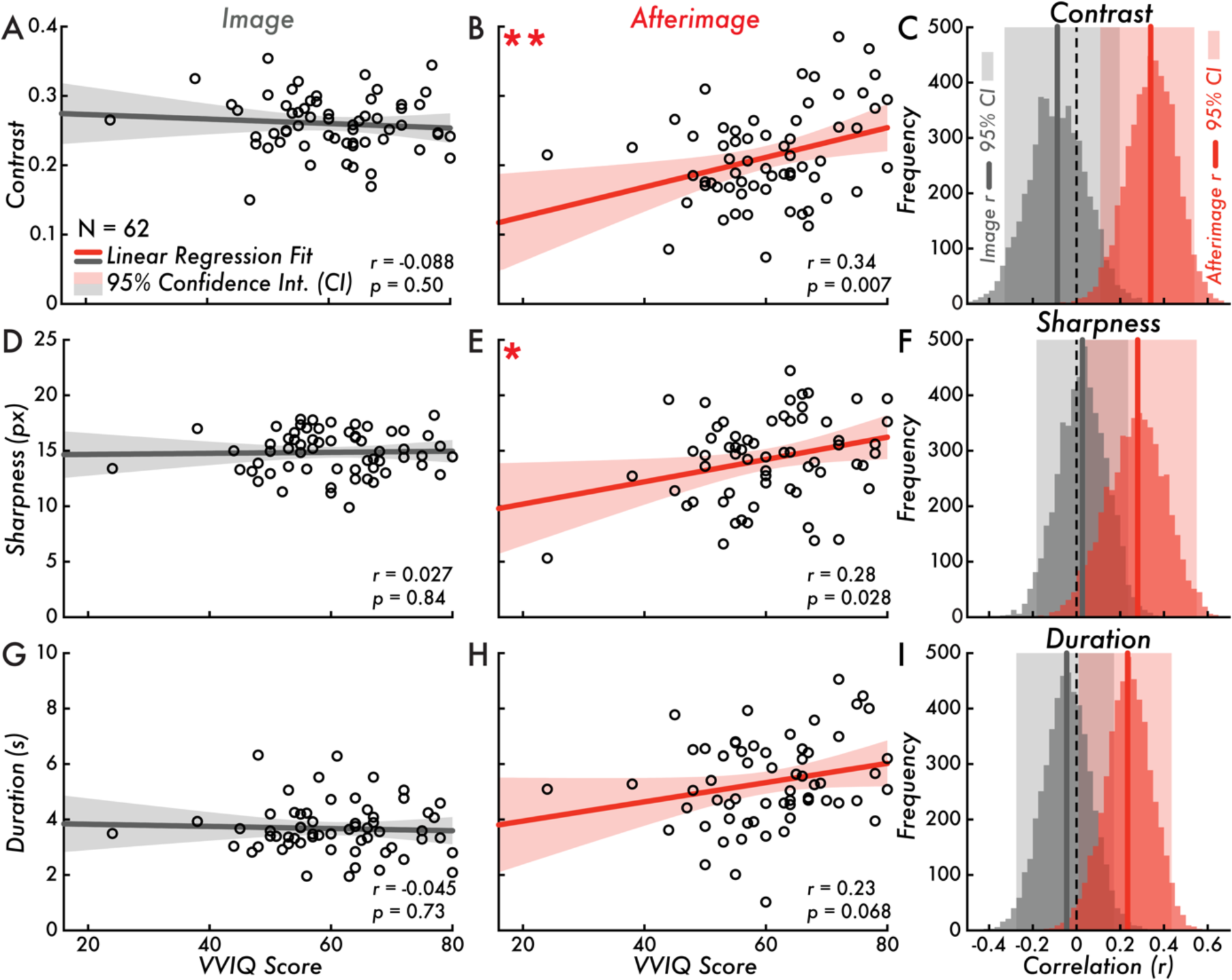
VVIQ score versus image and afterimage contrast, sharpness, and duration. **(A)** Vividness of Visual Imagery Questionnaire (VVIQ) score versus image contrast (correlation is not statistically significant; Pearson correlation coefficient [*r*] = −0.088; *p* = 0.50). **(B)** VVIQ score versus afterimage contrast (correlation is statistically significant **; *r* = 0.34; *p* = 0.007). **(C)** Bootstrapped image and afterimage VVIQ score and contrast correlation distributions and estimated 95% confidence interval (CI; image: [−0.33, 0.20]; afterimage: [0.11, 0.54]). **(D)** VVIQ score versus image sharpness (correlation is not statistically significant; *r* = 0.027; *p* = 0.84). **(E)** VVIQ score versus afterimage sharpness (correlation is statistically significant *; *r* = 0.28; *p* = 0.028). **(F)** Bootstrapped image and afterimage VVIQ score and sharpness correlation distributions and estimated 95% CI (image: [−0.18, 0.24]; afterimage: [0.041, 0.55]). **(G)** VVIQ score versus image duration (correlation is not statistically significant; *r* = −0.045; *p* = 0.73). **(H)** VVIQ score versus afterimage duration (correlation is not statistically significant; *r* = 0.23; *p* = 0.068). **(I)** Bootstrapped image and afterimage VVIQ score and duration correlation distributions and estimated 95% CI (image: [−0.28, 0.17]; afterimage: [0.01, 0.44]). Subplots A, B, D, E, G, and H, display the VVIQ score along the horizontal axis (score range: 16-80; larger values indicating more vivid visual imagery). The gray and red lines draw the linear regression fit of VVIQ score versus image or afterimage contrast, sharpness, and duration. The shaded area on either side of the main trend line is the 95% CI of the linear regression fit. The open circles represent individual participants (N = 62). In subplots C, F, and I, the gray and red vertical lines draw the Pearson correlation coefficient *r* value of VVIQ score versus image or afterimage contrast, sharpness, and duration. The shaded area behind the bootstrap correlation distributions is the estimated 95% CI.

**Figure 4.**
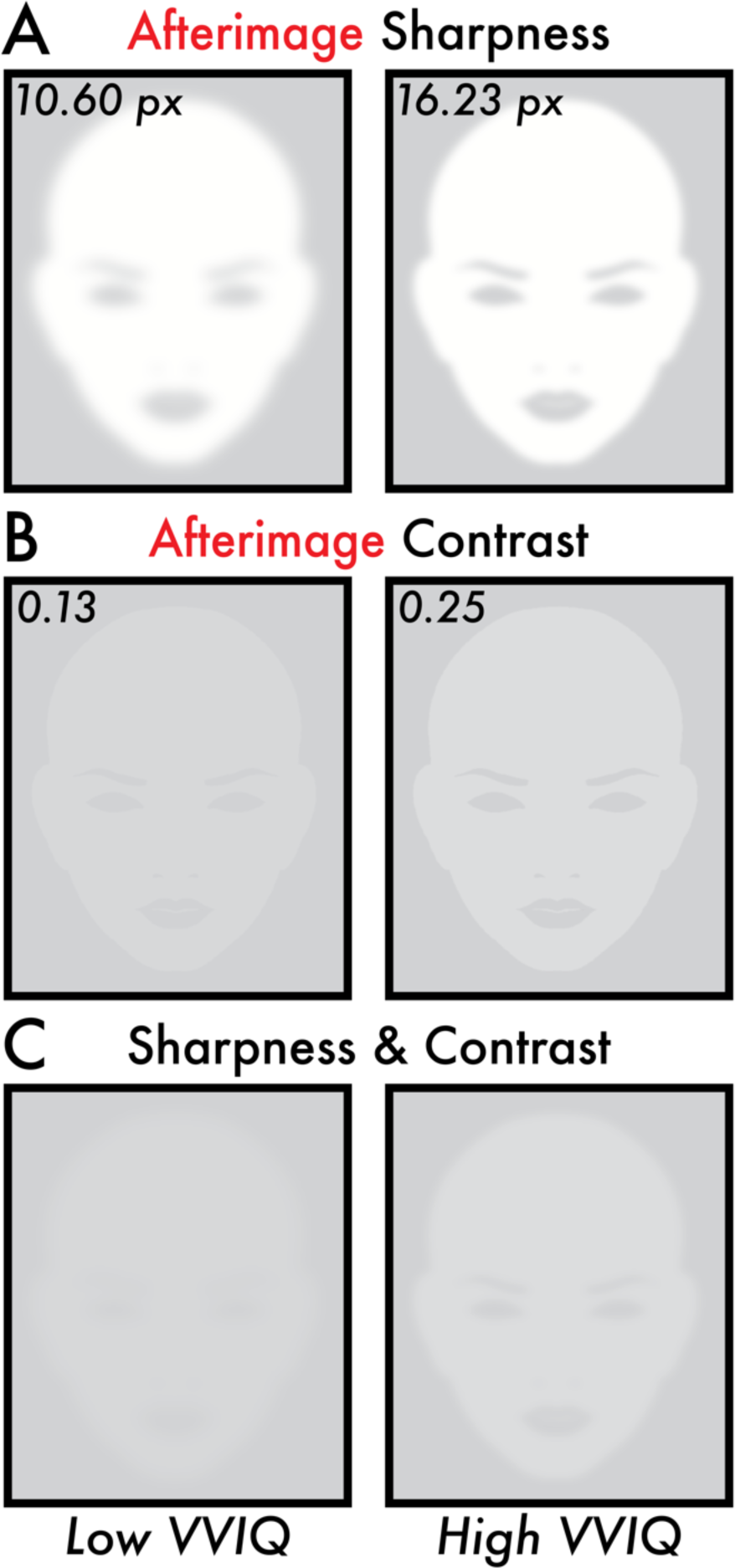
Image reconstruction of the estimated perceived afterimage sharpness and contrast for low and high VVIQ score participants. All subplots display image reconstructions of the estimated perceived afterimage sharpness and contrast values according to the fitted linear regression trend lines (Figure 3B, E) for low and high Vividness of Visual Imagery Questionnaire (VVIQ) scores. The low and high VVIQ scores are the minimum (24) and maximum (80) VVIQ scores reported among participants (Figure 2B). **(A)** Image reconstruction of the estimated perceived afterimage sharpness for low (10.60 pixels [px]) and high (16.23 px) VVIQ scores. **(B)** Image reconstruction of the estimated perceived afterimage contrast for low (0.13) and high (0.25) VVIQ scores. **(C)** Image reconstruction combining the estimated perceived afterimage sharpness (A) and contrast (B) for low (10.60 px and 0.13) and high (16.23 px and 0.25) VVIQ scores. Image reconstructions show apparent differences in overall visibility and facial feature details for the estimated perceived sharpness and contrast of afterimages between low and high VVIQ score participants.

Participants were instructed to report the maximum sharpness of the image stimulus in real time. This was achieved by the following steps within each image sharpness perception matching task trial (Supplementary Movie 2):

(1) Participants were instructed to fixate on a central plus sign inside an open circle (1.33 x 1.33 degrees) on a blank gray screen throughout the experiment (Figure 1D *Fixation* phase).
(2) After a jittered pre-stimulus interval (6-8 seconds), an image stimulus appeared (see image stimulus description above; Figure 1D *Image & Report* phase). The image would appear at random and in equal proportion either to the left or right of the fixation point along the midline (image stimulus location from central fixation = 5.88 degrees).
(3) Immediately upon perceiving the image stimulus, participants manually adjusted the sharpness of a controllable stimulus to match with the perceived maximum sharpness of the image stimulus using two keys: one increasing and the other decreasing the controllable image sharpness in increments of 1 pixel (Figure 1C, D *Controllable Images - Sharpness* and *Image & Report* phase). The controllable image was absent from the screen until the participant made their first key press to adjust its sharpness. The controllable image appeared at a random initial sharpness value (0-25 pixels) and was shown on the *opposite* side of the screen from where the image stimulus appeared.
(4) Once participants completed adjusting the sharpness of the controllable stimulus, they were instructed to press a third key to record their selection. While participants were encouraged to report the maximum sharpness of the image while the stimulus was still present on-screen, participants had a minimum of 10 seconds and a maximum of 12 seconds from the image offset (i.e., 10-12 seconds jittered post-image interval) to adjust the controllable image and make their perceived maximum sharpness selection (Figure 1D post-Image & Report *Fixation* phase). Otherwise, the trial was automatically aborted and no response was logged. A total of 20 trials of the image sharpness perception matching task was completed for each participant.

#### Afterimage Sharpness Perception Matching

Participants completed an *afterimage* sharpness perception matching task (Figure 1E; Supplementary Movie 3). The goal was for participants to report the *maximum* sharpness of their perceived afterimages. The reporting method and trial phases were identical to the image sharpness perception matching task (i.e., manually updating the sharpness of a controllable image with key presses to match with the perceived afterimage maximum sharpness; see *Image Sharpness Perception Matching* section). The main difference between the image and afterimage sharpness perception matching task phases was that in the afterimage condition, participants were first shown the inducer stimulus (see *Afterimage Induction* section; Figure 1E *Inducer* phase). When the inducer disappeared, the participants might see an afterimage on the blank gray screen and were instructed to immediately adjust the controllable image using the same reporting procedure as the image sharpness perception matching task (Figure 1E *Afterimage & Report* phase). If participants did not see an afterimage, they were instructed to *not* press any keys and wait until the next trial began automatically. A total of 30 trials of the afterimage sharpness matching task was completed for each participant.

### Contrast and Duration Perception Matching Paradigm

Participants were asked to report on the perceived contrast of images and afterimages *overtime* (i.e., to follow the change in the image and afterimage contrast throughout its perception). Previous studies have used perceptual cancellation to assess afterimage contrast (i.e., overlaying a physical image over the afterimage location and having participants adjust that physical image until the afterimage percept disappears; e.g., ^13^). In the current investigation, contrast was reported using a controllable, on-screen image that participants volitionally manipulated using key presses. See full details below.

#### Image Contrast and Duration Perception Matching

Participants completed an *image* contrast and duration perception matching task (Figure 1D; Supplementary Movie 4). The image was the same stimulus used in the image sharpness perception matching task (Figure 1C *Image Stimulus*). The image stimulus appeared for 4 seconds. In the first second of image presentation, the stimulus was shown gradually increasing its contrast to a maximum contrast of 0.25 (full contrast equals 1) and then gradually decreasing its contrast until the stimulus disappeared. The maximum contrast value (0.25) was selected according to pilot testing (data not shown) that suggested this contrast was similar to the maximum contrast of afterimages that appeared by the current inducer stimulus. There were three increasing contrast intervals (1, 1.5, and 2 seconds from the image stimulus onset until the image reached maximum contrast) and three decreasing contrast intervals (2.5, 3, and 3.5 seconds from the image stimulus onset until the image contrast began decreasing to full disappearance at 4 seconds). Thus, the image stimulus maintained the maximum contrast for a varied interval of 0.5, 1, 1.5, 2, and 2.5 seconds. The ramping contrast intervals were selected to approximate the contrast dynamic of the afterimage conscious perceptions reported in pilot testing (data not shown) from the current inducer stimulus.

Participants were instructed to report in real time the contrast of the image stimulus throughout its presentation. This was achieved by the following steps within each image contrast perception matching trial (Supplementary Movie 4):

(1) Participants were instructed to fixate on a central plus sign inside an open circle (1.33 x 1.33 degrees) on a blank gray screen throughout the experiment (Figure 1D *Fixation* phase).
(2) After a jittered pre-stimulus interval (6-8 seconds), an image stimulus appeared (see image stimulus description above; Figure 1D *Image & Report* phase). The image would appear at random and in equal proportion, either to the left or right of the fixation point along the midline (image stimulus location from central fixation = 5.88 degrees).
(3) Immediately upon perceiving the image stimulus, participants manually adjusted the contrast of a controllable image to match with the perceived contrast of the image stimulus overtime using two keys: one increasing and the other decreasing the controllable image contrast in increments of 0.025 (Figure 1C *Controllable Images - Contrast*). Participants could also use a third key that would immediately set the controllable image contrast to 0, thereby offering the option to report the perception of an immediate disappearance. Critically, participants were instructed to manipulate the controllable image to match with the image stimulus contrast *throughout* its presentation, so that at any given moment both the image and controllable image appeared with identical contrast. The controllable image appeared on the *opposite* side of the screen from the image stimulus.

The reported duration of the images was acquired by measuring the length of time participants manipulated the controllable image (i.e., the time when participants first reported a perceived image with greater than 0 contrast and its subsequent disappearance time; see the Statistical Analyses *Duration* section). While participants were encouraged to report the contrast of the image while the image stimulus was still present on-screen, participants had a minimum of 10 seconds and a maximum of 12 seconds from the image offset (i.e., 10-12 seconds jittered post-image interval) to continue adjusting the controllable image, after which the trial was automatically aborted and the responses made in the preceding interval were logged (Figure 1D post-Image & Report *Fixation* phase). A total of 18 trials of the image contrast and duration matching task was completed for each participant.

#### Afterimage Contrast and Duration Perception Matching

Participants completed an *afterimage* contrast and duration perception matching task (Figure 1E; Supplementary Movie 5). The goal was for participants to report the change in contrast overtime of their perceived afterimages. The reporting method and trial phases were identical to the image contrast and duration perception matching task (i.e., manually updating the contrast of a controllable image with key presses to match with the perceived afterimage contrast throughout its conscious perception; see *Image Contrast and Duration Perception Matching* section). The main difference between the image and afterimage contrast and duration perception matching task phases was that in the afterimage condition, participants were first shown the inducer stimulus (see *Afterimage Induction* section; Figure 1E *Inducer* phase). When the inducer disappeared, the participants might see an afterimage on the blank gray screen and were instructed to immediately display and adjust the controllable image using the same reporting procedure as the image contrast and duration perception matching task (Figure 1E *Afterimage & Report* phase). If participants did not see an afterimage, they were instructed to *not* press any keys and wait until the next trial began automatically. A total of 60 trials of the afterimage contrast and duration perception matching task was completed for each participant.

### Visual Imagery Vividness

Visual imagery vividness was acquired with the 16-item, self-reported Vividness of Visual Imagery Questionnaire (VVIQ)^40^. The questionnaire asks participants to imagine people, objects, and scenes (e.g., the contour of a familiar face, the front of a shop, and a sun rise) and then introspect on how vivid that imagined content appears in their visual imagery on a 5-point scale between “no image at all” to “perfectly clear and as vivid as normal vision”. Participants were instructed to complete the VVIQ with their eyes open and were given no time constraint in completing the questionnaire. The VVIQ was displayed on a computer monitor and participants used a mouse click to select their answers for each questionnaire item. The VVIQ was administered at either the beginning or end of the study session.

### Equipment, Software, and Facility

The behavioral study was completed in a single 2-hour study session in a windowless behavioral testing room. The room lighting was set to a consistent brightness level for all participants. The experimenter was present in the testing room but positioned out of sight of the participant to monitor behavior and deliver task instructions. The behavioral paradigm was coded in Python and run with PsychoPy (v2022.2.4; Open Science Tools Ltd.) on a behavioral laptop (MacBook Pro; 13-inch; 2560 x 1600 pixels, 2019; Mac OS Catalina v10.15.7; Apple, Inc.)^41^. The behavioral laptop monitor was mirrored by DVI cable to a VIEWPixx monitor (1920 x 1200 pixels; VPixx Technologies, Inc.) on which the participants viewed the experimental paradigms and the VVIQ. The participants were positioned approximately 56 cm from the center of the display monitor. The viewing distance was fixed using a table mounted head-chin rest. All participants used their right hand (regardless of handedness) to make key presses during the task with a keyboard positioned on a table in front of the participant.

## Statistical Analyses

All analyses were completed in MATLAB v2022b (MathWorks, Inc.) and Prism v10 (Graphpad, Inc.). Figures were generated and edited in MATLAB v2022b (MathWorks, Inc.), Prism v10 (Graphpad, Inc.), and Illustrator (Adobe, Inc.).

### Afterimage Perception Rate

Afterimage perception rate measures how often afterimages were perceived by each participant following the inducer stimulus. The perception rate was calculated by finding the percentage of inducer presentations that an afterimage was perceived across the sharpness and contrast and duration perception matching tasks – a total of 90 trials (i.e., the number of perceived afterimage trials in the sharpness perception matching task *plus* the number of perceived afterimage trials in the contrast and duration perception matching task *divided* by the total number of trials across all tasks). Perception rate values were multiplied by 100 to convert from units of fraction to percentage.

### VVIQ Score

The VVIQ score for each participant was calculated by taking the sum of all scores across the questionnaire items. Each item was scored on a scale from 1 (no image) to 5 (perfectly clear). Therefore, the minimum and maximum VVIQ score was 16 and 80, respectively, where larger values indicate more vivid visual imagery.

### Sharpness

#### Calculating reported sharpness

Participants reported the perceived maximum sharpness of images and afterimages (see *Sharpness Perception Matching Paradigm* Methods section). The participant image and afterimage sharpness values were calculated by averaging all trial sharpness values within participant and image and afterimage sharpness perception matching tasks. Trials without a sharpness value (e.g., response timeout or afterimage was not perceived) were excluded from consideration in calculating the participant maximum sharpness value. The sharpness value scale was inverted, so that larger values correspond with a sharper perception. This scale inversion was achieved by taking the *absolute value* of the participant mean sharpness value *minus* the maximum sharpness value (25; i.e., the largest pixel radius of the blurring gaussian kernel).

#### Calculating reported image sharpness accuracy

Participant reported image sharpness accuracy was calculated by subtracting the reported maximum image sharpness from the true image maximum sharpness (10, 15, or 20 pixels) across trials. Next, all subtracted or *relative* sharpness trial values were averaged within participant. A positive relative sharpness value indicated the image was reported as *sharper* than its true maximum sharpness, while a negative relative sharpness indicated the image was reported as *blurrier* than its true maximum sharpness, where a value of 0 indicated a perfect match between the reported and true image maximum sharpness (Figure 2C *Relative*). To statistically test the reporting accuracy of the image maximum sharpness, a Wilcoxon Rank Sum test (*p* < 0.05) was applied on the relative sharpness values and tested against 0. If the relative image sharpness is found no different from 0, then participants were accurate in reporting on the maximum sharpness of the image stimulus.

#### Calculating the correlation between VVIQ score and sharpness

The relationship between VVIQ scores and the reported image and afterimage maximum sharpness were statistically tested using a two-tailed, Pearson correlation test (*p* < 0.05; Figure 3D, E). Correlation analyses were applied in two comparisons: (1) VVIQ score versus *image* sharpness and (2) VVIQ score versus *afterimage* sharpness. A linear regression fit was applied to model the trend and 95% confidence interval for each of the comparisons. In addition, the bootstrap resampling method (5000 samples) estimated the 95% confidence interval for the correlation between VVIQ score and image and afterimage maximum sharpness (Figure 3F). A confidence interval that includes 0 suggests no correlation at the 5% significance level.

#### Image reconstruction of the afterimage sharpness

The perceived maximum sharpness of the afterimage was reconstructed for low (24) and high (80) VVIQ scores, representing the minimum and maximum VVIQ score recorded among participants (Figure 2B). Reconstruction was achieved by finding the sharpness value for the low and high VVIQ scores along the VVIQ score versus afterimage sharpness linear regression fit trend line and creating images (gaussian blur; Illustrator; Adobe, Inc.) that matched with these estimated sharpness values (Figure 4A, C).

### Contrast

#### Calculating reported contrast

Participants reported the perceived contrast of images and afterimages overtime (see *Contrast and Duration Perception Matching Paradigm* Methods section). The participant image and afterimage contrast values were calculated by finding the *maximum* contrast value reported in each image and afterimage contrast and duration perception matching task trial. Next, the maximum contrast values were averaged across trials within the image and afterimage conditions for each participant. Any trial with less than two reported contrast values or a maximum contrast value of 0 (e.g., an afterimage was not perceived) was ignored from calculating the participant image and afterimage contrast value.

#### Calculating reported image contrast accuracy

Participant reported image contrast accuracy was calculated by subtracting the reported maximum image contrast values from the known image maximum contrast value (0.25) across trials. Next, all subtracted or *relative* contrast trial values were averaged within participant. A positive relative contrast indicated the image was reported as *brighter* than its true maximum contrast, while a negative relative contrast indicated the image was reported as *dimmer* than its true maximum contrast, where a value of 0 indicated a perfect match between the participant reports and the true image maximum contrast (Figure 2D *Relative*). To statistically test the reporting accuracy of the image maximum contrast, a Wilcoxon Rank Sum test (*p* < 0.05) was applied on the relative contrast values and tested against 0. If the relative image contrast is found no different from 0, then participants were accurate in reporting on the contrast of the image stimulus.

#### Calculating the correlation between VVIQ score and contrast

The relationship between VVIQ scores and the reported image and afterimage maximum contrast were statistically tested using a two-tailed, Pearson correlation test (*p* < 0.05; Figure 3A, B). Correlation analyses were applied in two comparisons: (1) VVIQ score versus *image* contrast and (2) VVIQ score versus *afterimage* contrast. A linear regression fit was applied to model the trend and 95% confidence interval for each of the comparisons. In addition, the bootstrap resampling method (5000 samples) estimated the 95% confidence interval for the correlation between VVIQ score and image and afterimage maximum contrast (Figure 3C). A confidence interval that includes 0 suggests no correlation at the 5% significance level.

#### Image reconstruction of the afterimage contrast

The perceived maximum contrast of the afterimage was reconstructed for low (24) and high (80) VVIQ scores, representing the minimum and maximum VVIQ score recorded among participants (Figure 2B). Reconstruction was achieved by finding the contrast value for the low and high VVIQ scores along the VVIQ score versus afterimage contrast linear regression fit trend line and creating images (Illustrator; Adobe, Inc.) that matched with these estimated contrast values (Figure 4B, C).

### Duration

#### Calculating reported duration

Image and afterimage durations were calculated from the contrast and duration perception matching tasks (see *Contrast and Duration Perception Matching Paradigm* Methods section). Contrast and duration perception matching task trials were considered valid by the same criteria for calculating the reported maximum contrast of images and afterimages (see *Contrast* Statistical Analyses section). The duration was measured as the time between the initial and final key press participants made to adjust the controllable image to match with the perceived contrast of the images and afterimages or when the participant reported the image or afterimage had a contrast of zero, whichever occurred first.

#### Calculating reported image duration accuracy

Participant reported image duration accuracy was calculated by subtracting the reported image duration across trials within participant from the true image duration (4 seconds). A positive relative duration indicated participants reported on average that the image was presented *longer* than its true duration, while a negative relative duration indicated that participants reported on average that the image was *briefer*, where a value of 0 indicated a perfect match between the reported and true image duration (Figure 2E *Relative*). To statistically test how accurate participants were in reporting the image duration, a Wilcoxon Rank Sum test (*p* < 0.05) was applied on the relative duration values and tested against 0. If the relative image duration is found no different from 0, then participants were accurate in reporting on the duration of the image stimulus.

#### Calculating the correlation between VVIQ score and duration

The relationship between VVIQ scores and the reported image and afterimage duration were statistically tested using a two-tailed, Pearson correlation test (*p* < 0.05; Figure 3G, H). Correlation analyses were applied in two comparisons: (1) VVIQ score versus *image* duration and (2) VVIQ score versus *afterimage* duration. A linear regression fit was applied to model the trend and 95% confidence interval for each of the comparisons. In addition, the bootstrap resampling method (5000 samples) estimated the 95% confidence interval for the correlation between VVIQ score and image and afterimage duration (Figure 3I). A confidence interval that includes 0 suggests no correlation at the 5% significance level.

## Results

### Afterimage Perception Rate and VVIQ Score

The inducer stimulus consistently induced afterimages in most participants: a mean afterimage perception rate of 90.79% (standard deviation [SD] = 14.84%; minimum participant afterimage perception rate = 38.89%; maximum participant afterimage perception rate = 100%; Figure 2A). The mean VVIQ score was 60.55 (SD = 10.78; minimum participant VVIQ score = 24; maximum participant VVIQ score = 80; Figure 2B). One participant was below the ∼30-score threshold that is commonly used to designate *aphantasia* – the near or total inability to form visual imagery – estimated to account for less than 5% of the general population^42^.

### Image and Afterimage Perceptual Features

Participants reported on their perceived sharpness, contrast, and duration of images and afterimages. The mean image and afterimage maximum sharpness values were 14.87 pixels (SD = 1.91 pixels) and 14.27 pixels (SD = 3.88 pixels), respectively (Figure 2C *Observed*; image observed not shown). The mean image and afterimage maximum contrast values were 0.26 (SD = 0.04) and 0.21 (SD = 0.068), respectively (Figure 2D *Observed*; image observed not shown). The mean image and afterimage duration values were 3.67 seconds (SD = 0.93 seconds) and 5.35 seconds (SD = 1.60 seconds), respectively (Figure 2E *Observed*; image observed not shown). These results revealed broad individual variability for the perceived sharpness, contrast, and duration of afterimages.

### Accuracy of the Reported Image Sharpness, Contrast, and Duration

A key validation was whether participants could accurately report on their perceptual experiences using the current perception matching paradigms (see *Sharpness Perception Matching Paradigm* and *Contrast and Duration Perception Matching Paradigm* Methods sections). Image stimuli with known sharpness, contrast, and duration were used to validate the accuracy of the perceptual reports using the perception matching paradigms. In support of the perception matching paradigms and their resulting perceptual reports, there was *no* statistically significant difference (*p* > 0.05) between the reported and true image sharpness, contrast, and duration. Specifically, the reported sharpness, contrast, and duration *minus* the true image sharpness, contrast, and duration were not statistically different from 0 (i.e., a perfect match between the reported and true image sharpness, contrast, and duration; Figure 2C, D, E *Relative*; see *Methods* section for image features and statistical testing details).

### VVIQ Score Versus Image Sharpness, Contrast, and Duration

Task-based factors, including motor and cognitive processes involved in reporting on the perceptual features of conscious perception could explain a relationship between VVIQ score and reported image and afterimage sharpness, contrast, and duration. This alternative hypothesis was tested by comparing VVIQ score with the reported *image* sharpness, contrast, and duration. There was *no* statistically significant correlation for VVIQ score versus image contrast (Pearson correlation coefficient [*r*] = - 0.088; *p* = 0.50; Figure 3A), VVIQ score versus image sharpness (*r* = 0.027; *p* = 0.84; Figure 3D), and VVIQ score versus image duration (*r* = −0.045; *p* = 0.73; Figure 3G). The bootstrap estimated 95% confidence intervals of the correlation distributions confirmed that there was no relationship between VVIQ score and image sharpness ([−0.18, 0.24]; Figure 3F), contrast ([−0.33, 0.20]; Figure 3C), and duration ([−0.28, 0.17]; Figure 3I).

### VVIQ Score Versus Afterimage Sharpness, Contrast, and Duration

To test for a link between the vividness of visual imagery and afterimages, VVIQ scores were correlated with the afterimage sharpness, contrast, and duration. A statistically significant, moderate positive correlation was found between VVIQ score and afterimage contrast (*r* = 0.34; *p* = 0.007; linear regression fit trend line equation: Y = 0.0021*X + 0.083; Figure 3B). The estimated contrast value according to the linear regression fit trend line for a low VVIQ score (minimum participant VVIQ score = 24) was 0.13 and a high VVIQ score (maximum participant VVIQ score = 80) was 0.25 (Figure 4B, C). A statistically significant, weak positive correlation was found between VVIQ score and afterimage sharpness (*r* = 0.28; *p* = 0.028; linear regression fit trend line equation: Y = 0.10*X + 8.19; Figure 3E). The estimated sharpness value according to the linear regression fit trend line for a low VVIQ score (minimum participant VVIQ score = 24) was 10.60 pixels and a high VVIQ score (maximum participant VVIQ score = 80) was 16.23 pixels (Figure 4A, C). There was no statistically significant correlation between VVIQ score and afterimage duration (*r* = 0.23; *p* = 0.068; Figure 3H). The bootstrap estimated 95% confidence intervals of the correlation distributions confirmed a positive correlation between VVIQ score and afterimage contrast ([0.11, 0.54]; Figure 3C) and sharpness ([0.041, 0.55]; Figure 3F), and supported a positive correlation between VVIQ score and afterimage duration ([0.01, 0.44]; Figure 3I).

When the recruited participant with aphantasia (see *Afterimage Perception Rate and VVIQ Score* section) is removed from the data set, the correlation between VVIQ score and afterimage contrast and duration become *stronger* (contrast: *r* = 0.38; *p* = 0.0026; duration: *r* = 0.25; *p* = 0.052). Meanwhile, because this participant reported the *least* sharp (i.e., blurriest) afterimages of all participants, when removed from analyses the correlation between VVIQ score and afterimage sharpness maintained the same trend but was no longer statistically significant (*r* = 0.17; *p* = 0.18). These results support a statistically robust correlation between VVIQ score and afterimage contrast. Meanwhile, additional research is required (e.g., testing more participants with aphantasia) to confirm a possible relationship between VVIQ score and afterimage sharpness and duration.

## Discussion

The current investigation is the first study relating the perception of imagery and afterimages. Our main finding was evidence for a perceptual link between visual imagery and negative afterimages. To interrogate this relationship, we developed novel perception matching paradigms where participants manipulated the appearance of on-screen images to report their perceived afterimage sharpness, contrast, and duration. The efficacy of these reporting methods was validated by testing the participant reporting accuracy for image stimuli with known sharpness, contrast, and duration (Figures 2C, D, E *Relative*). These perceptual reports revealed variability in the perceived sharpness, contrast, and duration of afterimages across participants (Figures 2C, D, E *Observed*). Moreover, we discovered a statistically significant, moderate positive correlation between visual imagery vividness and the perceived afterimage contrast (Figure 3B, C). In short, people who reported more vivid visual imagery tended to report brighter afterimages. There was also a weak positive correlation between visual imagery vividness and afterimage sharpness, however, this statistical result was sensitive to the participant with aphantasia (Figure 3E, F; see *Afterimage Perception Rate and VVIQ Score* Results section).

Bolstering these results was the specificity of the relationship of visual imagery vividness to afterimage conscious perception. Meanwhile, no correlation was found between VVIQ score and *image* contrast, sharpness, and duration in the same participants (Figure 3A, C, D, F, G, I). Moreover, reconstructions of the estimated afterimage conscious perceptions for low and high VVIQ scores were visibly distinct – the high VVIQ score afterimage reconstruction revealed an apparently brighter image with sufficient sharpness to discern facial features that were absent in the low VVIQ afterimage reconstruction (Figure 4C *Low* versus *High VVIQ*).

The relationship between the vividness of visual imagery and afterimage conscious perception is a novel source of behavioral evidence that imagery and afterimages may share neural mechanisms. Specifically, the afterimages induced in the current investigation may involve top-down, brain mechanisms in common with visual imagery that helps explain the observed individual variability in the perceived afterimage sharpness, contrast, and duration. This interpretation of the current results is corroborated by observations that afterimages can form without bottom-up, retinal stimulation (e.g., ^23,30^). However, these behavioral findings can only speculate on neural mechanisms. Thereby, our results encourage combining afterimage paradigms with neuroimaging and other neurophysiology recording methods – these studies currently limited in the literature – to directly examine the neural mechanisms of afterimage conscious perception.

An alternative explanation of these findings is the influence of mediating sensory, cognitive, or behavioral variables required by the perception reporting tasks (e.g., reaction time and sensory sensitivity) and reporting on visual imagery vividness. This account is dampened because reporting sharpness, contrast, and duration of images and afterimages versus visual imagery vividness involved orthogonal tasks (see *Image and Afterimage Perceptual Vividness* Methods section). Specifically, reporting on image and afterimage conscious perceptions required adjusting a controllable image with key presses. Meanwhile, visual imagery vividness was inquired using a self-paced questionnaire (i.e., the VVIQ) that involved marking responses with a mouse click. Therefore, sensory, cognitive, and behavioral ability are excluded as likely factors influencing the current findings. Indeed, if mediating variables linked to the task requirements explained these results, we would also expect a relationship between the vividness of visual imagery and *image* conscious perception, as identical perception reporting procedures were involved for both images and afterimages. Thus, perceptual reporting itself is unlikely to explain the current findings. Still, we cannot rule out the influence of other unknown factors that may be shared across reporting methods and unique to afterimages (e.g., metacognitive or introspection ability).

A challenge for the current investigation is assuring accuracy of the afterimage perceptual reports. Previous studies have manipulated the inducer stimulus to predictably change afterimage conscious perception to determine the veracity of perceptual reports^13^. Here, we used images with known sharpness, contrast, and duration to validate reporting on the same perceptual features of afterimages. In support of the perception reporting methods and reporting accuracy, participants were accurate for indicating the sharpness, contrast, and duration of images. Importantly, the image stimuli were designed to approximate the perceptual features of the afterimages to match the reporting difficultly between images and afterimages. Still, reporting on-screen images versus illusory afterimages may be distinct (e.g., more challenging to report on illusory conscious perceptions). Therefore, it is possible that image reporting accuracy may not predict afterimage reporting accuracy. Future studies may address this limitation by recording physiological markers of conscious perception (e.g., pupil size) that may serve as a covert measure of afterimage perceptual vividness to corroborate the veracity of overt perceptual reports. Likewise, pupil size has been shown to validate self-report of visual imagery vividness^43^.

The current findings motivate future study on the relationship between afterimages and other categories of sensory-independent conscious perception (e.g., hallucinations and dreams) in healthy physiology. Furthermore, it would be valuable to research how afterimages may change in psychiatric and neurologic disorders that impact sensory and sensory-independent conscious perception. Supporting this research aim, a study found differences in afterimage onset latency among people with brain injury^44^. In addition, afterimages are altered in people with autistic traits and schizophrenia^45–47^. These findings suggest afterimages may offer translational value, as previously suggested in posterior cortical atrophy – a variant of Alzheimer’s disease – and Parkinson’s disease where afterimages have also been found to be modified relative to healthy individuals^48–50^.

## Conclusion

Afterimages have long been a source of curiosity and implemented as a perceptual tool to interrogate vision and the neural mechanism of consciousness. In the current investigation, we studied a possible link between visual imagery and afterimage conscious perception. We developed novel perception matching paradigms that allowed for the acquisition of various perceptual features of image and afterimage conscious perception. Our main result was a correlation between the perceived vividness of visual imagery and negative afterimages. This represents a new source of evidence that some afterimages may share neural mechanisms with visual imagery. This study motives future research to directly examine the precise neural mechanisms of afterimage conscious perception.

## Author Contributions

S.I.K. contributed to conceptualization, methodology, software, formal analysis, investigation, data curation, visualization, supervision, project administration, and writing (original draft); M.H. contributed to methodology, software, investigation, and writing (review and editing); A.T.M. contributed to conceptualization, methodology, and writing (review and editing); J.B.T. contributed to methodology, software, and writing (review and editing); J.G-C. contributed to formal analysis and writing (review and editing); D.A.H. contributed to conceptualization, methodology, supervision, formal analysis, and writing (review and editing); P.A.B. contributed to conceptualization, methodology, supervision, project administration, funding acquisition, and writing (review and editing).

## Acknowledgements

This research was made possible by the support of the National Institute of Mental Health Intramural Research Program (ZIAMH002783). The study was completed in compliance with the National Institutes of Health Clinical Center protocol ID 93-M-0170 (ClinicalTrials.gov ID: NCT00001360). We thank members of the Section on Functional Imaging Methods for their constructive feedback.

## Data and Code Availability

All data and scripts will be made available prior to publication at https://github.com/nimh-sfim

