## Supplementary Information for "Visual imagery vividness correlates with afterimage conscious perception"

**Contents:**

*Supplementary Movie 1 Legend:* Inducer stimulus
*Supplementary Movie 2 Legend:* Image sharpness perception matching
*Supplementary Movie 3 Legend:* Afterimage sharpness perception matching
*Supplementary Movie 4 Legend:* Image contrast and duration perception matching
*Supplementary Movie 5 Legend:* Afterimage contrast and duration perception matching

**Supplementary Movie 1.** *Inducer stimulus and afterimage demonstration.* Initially a central fixation point (a plus sign inside an open circle) appears on a blank grey background followed by the presentation of the inducer stimulus (see *Afterimage Induction* Methods section; Figure 1C *Inducer Stimulus*). After the inducer stimulus disappears, a negative afterimage (i.e., a light grey or white, blurrier version of the inducer) may become visible after several seconds in the same location that the inducer stimulus was shown, if maintaining fixation throughout the demonstration. A minority of individuals may not experience an afterimage.

**Supplementary Movie 2.** *Image sharpness perception matching task trial.* In the image sharpness perception matching task, participants reported the *maximum* sharpness of image stimuli by manipulating the sharpness of a controllable image (see *Image Sharpness Perception Matching* Methods section; Figure 1C *Image Stimulus* and *Controllable Images - Sharpness*, D). This movie clip demonstrates a single trial of the image sharpness matching procedure. Initially a central fixation point (a plus sign inside an open circle) appears on a blank gray background. Next, an image stimulus appears on the left side of the fixation point. The image stimulus subtly changes its sharpness overtime before disappearing from the screen. Simultaneously, the participant manually presents the controllable image on the right side of the fixation point and adjusts the sharpness of the controllable image using key presses to match with the perceived maximum sharpness of the image stimulus, all while maintaining central fixation. When the participant has satisfactorily matched the sharpness of the controllable image to the maximum sharpness of the preceding image stimulus, they press the selection key that logs their response and immediately removes the controllable image from the screen.

**Supplementary Movie 3.** *Afterimage sharpness perception matching task trial.* In the afterimage sharpness perception matching task, participants reported the *maximum* sharpness of their afterimages by manipulating the sharpness of a controllable image (see *Afterimage Sharpness Perception Matching* Methods section; Figure 1C *Controllable Images - Sharpness*, E). This movie clip demonstrates a single trial of the afterimage sharpness matching procedure. Initially a central fixation point (a plus sign inside an open circle) appears on a blank gray background. Next, an inducer stimulus appears on the right side of the fixation point (Figure 1C *Inducer Stimulus*). After the inducer stimulus disappears, the participant perceives an afterimage on the left side of the fixation point, although nothing physically appears on screen, and manually presents the controllable image on the right side of the fixation point. The participant adjusts the sharpness of the controllable image using key presses to match with the perceived maximum sharpness of their afterimage, all while maintaining central fixation. When the participant has satisfactorily matched the sharpness of the controllable image to the maximum sharpness of their afterimage, they press the selection key that logs their response and immediately removes the controllable image from the screen.

**Supplementary Movie 4.** *Image contrast and duration perception matching task trial.* In the image contrast and duration perception matching task, participants reported the contrast overtime of image stimuli by manipulating the contrast of a controllable image (see *Image Contrast and Duration Perception Matching* Methods section; Figure 1C *Image Stimulus* and *Controllable Images - Contrast*, D). This movie clip demonstrates a single trial of the image contrast and duration matching procedure. Initially a central fixation point (a plus sign inside an open circle) appears on a blank gray background. Next, an image stimulus appears on the left side of the fixation point. The image stimulus subtly changes its contrast overtime before disappearing from the screen. Simultaneously, the participant manually presents the controllable image on the right side of the fixation point and adjusts the contrast of the controllable image using key presses to match with the perceived contrast of the image stimulus overtime, all while maintaining central fixation. The participant continues this matching procedure until the image stimulus disappears and concurrently removes the controllable image from the screen.

**Supplementary Movie 5.** *Afterimage contrast and duration perception matching task trial.* In the afterimage contrast and duration perception matching task, participants reported the contrast overtime of their afterimages by manipulating the contrast of a controllable image (see *Afterimage Contrast and Duration Perception Matching* Methods section; Figure 1C *Controllable Images - Contrast*, E). This movie clip demonstrates a single trial of the afterimage contrast and duration matching procedure. Initially a central fixation point (a plus sign inside an open circle) appears on a blank gray background. Next, an inducer stimulus appears on the left side of the fixation point (Figure 1C *Inducer Stimulus*). After the inducer stimulus disappears, the participant perceives an afterimage on the left side of the fixation point, although nothing physically appears on screen, and manually presents the controllable image on the right side of the fixation point. The participant adjusts the contrast of the controllable image using key presses to match with the perceived contrast of their afterimage overtime, all while maintaining central fixation. The participant continues this matching procedure until their afterimage perception disappears and concurrently removes the controllable image from the screen.
